# Gene networks driven by genetic variation for plasma cortisol in hepatic and adipose tissues implicate corticosteroid binding globulin in modulating tissue glucocorticoid action and cardiovascular risk

**DOI:** 10.1101/2023.01.20.524857

**Authors:** Sean Bankier, Lingfei Wang, Andrew Crawford, Ruth A Morgan, Arno Ruusalepp, Ruth Andrew, Johan LM Björkegren, Brian R Walker, Tom Michoel

**Affiliations:** University/ BHF Centre for Cardiovascular Science, Queen’s Medical Research Institute, University of Edinburgh, Edinburgh, UK; Computational Biology Unit, Department of Informatics, University of Bergen, PO Box 7803, 5020 Bergen, Norway; Division of Genetics and Genomics, The Roslin Institute, The University of Edinburgh, Easter Bush, Midlothian, UK; MRC Integrative Epidemiology Unit, University of Bristol, Bristol, UK; SRUC, The Roslin Institute, Easter Bush, Edinburgh, UK; Department of Cardiac Surgery, Tartu University Hospital, Tartu, Estonia; Department of Cardiology, Institute of Clinical Medicine, Tartu University, Tartu, Estonia; Clinical Gene Networks AB, Stockholm, Sweden; Department of Medicine, Karolinska Institutet, Karolinska Universitetssjukhuset, Huddinge, Sweden; Department of Genetics & Genomic Sciences, Institute of Genomics and Multiscale Biology, Icahn School of Medicine at Mount Sinai, New York, NY, USA; Clinical and Translational Research Institute, Newcastle University, Newcastle upon Tyne, UK

## Abstract

Genome wide association meta-analyses (GWAMA) by the CORtisol NETwork (CORNET) consortium identified genetic variants spanning the *SERPINA6/ SERPINA1* locus on chromosome 14 associated with morning plasma cortisol, cardiovascular disease (CVD), and *SERPINA6* mRNA expression encoding corticosteroid binding globulin (CBG) in liver. These and other findings indicate that higher plasma cortisol levels are causally associated with cardiovascular disease, however, the mechanisms by which variations in CBG lead to CVD are undetermined. Using genomic and transcriptomic data from The Stockholm Tartu Atherosclerosis Reverse Networks Engineering Task (STARNET) study, we identified plasma cortisol linked Single Nucleotide Polymorphisms (SNPs) that are trans-associated with genes from 7 different vascular and metabolic tissues, finding the highest representation of transgenes in liver, subcutaneous adipose and visceral abdominal adipose tissue (FDR = 15%). We identified a sub-set of cortisol-associated trans-genes that are putatively regulated by the Glucocorticoid Receptor (GR), the primary transcription factor activated by cortisol. Using causal inference, we identified GR-regulated trans-genes that are responsible for the regulation of tissue specific gene networks. Cis-expression Quantitative Trait Loci (eQTLs) were used as genetic instruments for identification of pairwise causal relationships from which gene networks could be reconstructed. Gene networks were identified in liver, subcutaneous fat and visceral abdominal fat, including a high confidence gene network specific to subcutaneous adipose (FDR = 10%) under the regulation of the interferon regulatory transcription factor, *IRF2*. These data identify a plausible pathway through which variation in liver CBG production perturbs cortisol-regulated gene networks in peripheral tissues and thereby promote CVD.

## Introduction

The steroid cortisol is the major glucocorticoid hormone involved in mediating the human stress response, with effects on metabolism, cardiovascular homeostasis and inflammation^1^. Excessive cortisol production occurs in Cushing’s syndrome, either in response to chronic activation of the hypothalamic-pituitary-adrenal (HPA) axis by increased Adrenocorticotropic Hormone (ACTH) secretion or through autonomous production of cortisol in an adrenocortical tumour^2^. Cushing’s syndrome results in insulin resistance, obesity and hypertension with increased risk of CVD. Similarly, higher plasma cortisol within the population, in the absence of overt Cushing’s syndrome, is associated with risk factors for cardiovascular disease (CVD) such as hypertension^3^ and type II diabetes^1,4^.

Inter-individual variation in plasma cortisol levels has a genetic basis with heritability estimated between 30-60%^5^. The CORtisol NETwork (CORNET) consortium conducted a Genome Wide association Meta-Analysis (GWAMA) with the intention of uncovering genetic influences on HPA axis function^6^. This was followed in 2021 with an expanded GWAMA of 25,314 individuals across 17 population-based cohorts of European ancestries^7^. In an additive genetic model, the CORNET GWAMA identified 73 genome-wide significant Single Nucleotide Polymorphisms (SNPs) associated with variation for plasma cortisol at a single locus on chromosome 14. These SNPs were used in a two-sample Mendelian Randomization analysis showing that higher cortisol is causative for CVD^7^.

The locus on chromosome 14 spans the genes *SERPINA6* and *SERPINA1* which both play roles in the regulation of Corticosteroid Binding Globulin (CBG), a plasma protein produced in liver which is responsible for binding 80-90% of cortisol in the blood^8,9^. *SERPINA6* encodes CBG10 and *SERPINA1* encodes α1-antitrypsin, an inhibitor of neutrophil elastase, a serine protease which can cleave the reactive centre loop of CBG resulting in a 9-10 fold reduction in binding affinity to cortisol^11,12^.

The CORNET GWAMA showed that 21 cortisol-associated SNPs were also cis-expression Quantitative Trait Loci (eQTLs) for *SERPINA6* in liver and demonstrated that the genetic variation associated with plasma cortisol is driven by *SERPINA6* rather than *SERPINA1^7^*. However, although variation in CBG production could explain changes in total plasma cortisol, it is the free fraction of cortisol that is considered to equilibrate with target tissue concentrations and signal through intracellular glucocorticoid receptors (GR). While CBG deficiency may be associated with symptoms^13–15^, variations in CBG have not been shown conclusively to influence the tissue response to cortisol in humans.

To test the hypothesis that cortisol-associated genetic variants in the *SERPINA6/SERPINA1* locus influence cortisol delivery to, and hence action in, extra-hepatic tissues, we investigated transcriptome-wide associations between cortisol-associated SNPs and gene transcripts across 7 different vascular and metabolic tissues from the Stockholm Tartu Atherosclerosis Reverse Networks Engineering Task study (STARNET) study^16^. As well as conducting a multi-tissue eQTL analysis using STARNET transcriptomics and plasma cortisol-associated SNPs, we identified tissuespecific trans-eQTL-associated genes under the regulation of GR. Moreover, we used a causal inference framework, with cis-eQTLs as genetic instruments, for the reconstruction of causal gene networks within STARNET tissues.

The results provide evidence that genetic variations in CBG production in liver influence extra-hepatic cortisol signaling and provide plausible pathways leading to CVD.

## 2 Materials and methods

### 2.1 Data

STARNET is a cohort-based study of 600 individuals undergoing Coronary Artery Bypass grafting (CABG) for Coronary Artery Disease (CAD) and was used as the primary discovery cohort in this study. These individuals underwent blood genotyping pre-operatively and during surgery. 7 different tissue samples were obtained and underwent RNA-sequencing: liver, skeletal muscle, atherosclerotic aortic root, internal mammary artery, visceral abdominal fat, subcutaneous fat, and whole blood. STARNET data are available through a database of Genotypes and Phenotypes (dbGaP) application (accession no. phs001203.v2.p1).

The Stockholm Atherosclerosis Gene Expression study (STAGE) (n=114)^17^ and the Metabolic Syndrome in Man study (METSIM) (n=982)^18^ were used in the replication of causal gene networks identified using STARNET. Gene expression data for METSIM and STAGE are available publicly at GEO (accession no. GSE70353 and GSE40231, respectively). Microarray data for liver, subcutaneous fat and visceral abdominal fat was used from STAGE and gene expression data from subcutaneous fat was measured in METSIM using RNA-sequencing (RNA-seq).

### 2.2 Processing of genotype and tissue-specific gene expression data

All STARNET genotype and gene expression data obtained for this project had undergone both Quality Control (QC) and normalisation as described previously^16^. The Human OmniExpressExome-8v1 bead chip was used with GRCh37 and contains 951,117 genomic markers. Genotypes were contained in matrices within the −012 format and a filtering step was included to remove any SNPs which had missing values for any samples and to exclude any SNPs which had a Minor Allele Frequency (MAF) < 5%.

Gene expression for STARNET tissue samples was measured using RNA-seq. RNA samples with less than 1M uniquely mapped reads were excluded, which removed 12 samples with extremely low read counts. The read counts of the samples used in the final analysis were between 15-30 million reads (Figure S1).

The numbers of samples and genes retained can be seen in Table 1. Having obtained gene expression matrices from Franzen *et al*, we conducted Principal Component Analysis (PCA) to confirm that there were no outliers within the samples (Figure S2). Ensembl Biomart (GRCh37) was used to label transcripts (provided as Ensembl IDs) with gene name, chromosome location, gene start, gene end and the Transcription Start Site (TSS).

**Table 1:** Summary of tissue specific gene expression data from STARNET.

| Tissue | Number of samples | Number of genes per sample |
| --- | --- | --- |
| Liver | 523 | 13875 |
| Skeletal muscle | 512 | 12544 |
| Internal mammary artery | 529 | 15458 |
| Atherosclerotic aortic root | 514 | 16214 |
| Subcutaneous fat | 549 | 14120 |
| Visceral abdominal fat | 509 | 14965 |
| Whole blood | 448 | 12843 |

METSIM gene expression data were obtained as Transcripts Per Million (TPM) and showed an inflation of higher correlated genes from the normal distribution (Figure S4). To account for this, the METSIM gene expression values were log2 transformed (TPM+1) followed by a rerunning of the PCA. The log2 transformed expression values were then fitted to a linear model in R, while adjusting on the 1st principal component. The residuals of this model replaced the count values that were used in all subsequent analyses and no longer showed inflated correlation values (Figure S4).

### 2.3 Multi-tissue trans-eQTL discovery

A list of SNPs associated with plasma cortisol was obtained from the summary statistics of the 2021 GWAMA conducted by the CORNET consortium (available at https://datashare.ed.ac.uk/handle/10283/3836)^7^. We filtered this list to obtain SNPs that were found to be associated with plasma cortisol at a level of genome wide significance (p ≤ 5×10-8) which were taken forward and tested against all genes across STARNET tissues.

The secondary linkage test (P2) is a likelihood ratio test in the Findr package^19^ (version 1.0.8) that was used to identify associations between a given SNP (E) and a gene (B) using categorical regression. P2 proposes a null hypothesis where E and B are independent and alternative hypothesis where E is causal for B (E→B). Maximum likelihood estimators are then used to obtain a log likelihood ratio (LLR) between the alternative and null hypothesis. The LLR is then converted to the posterior probability of the alternative hypothesis 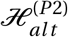 being true with empirical estimation of the local False Discovery Rate (FDR) as a value from 0-1 (Equation 1).

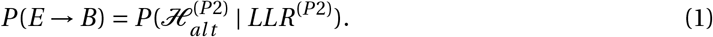

### 2.4 Identification of glucocorticoid-regulated trans-genes

Multiple datasets were used to identify genes that had prior evidence of regulation by GR (Table 2). These datasets have been filtered to include targets for *NR3C1*, the gene which encodes GR.

**Table 2:** Datasets used for identifying genes regulated by glucocorticoids.

| Dataset | Study type | Target definition |
| --- | --- | --- |
| Yu *et al*, 2010<sup>20</sup> | ChIP-seq and microarray analysis in adipocyte specific (3T3-L1). | Use of PinkThing to identify gene closest to peak. |
| Bell *et al*, 2021<sup>21</sup> | RNA-seq for subcutaneous and epididymal adipose from dexamethasone treated mice. | Differential expression based on fold change. |
| ENCODE<sup>22</sup> | Large scale high throughput ChIP-seq. | Map ChIP-seq peak to closest gene. If multiple peaks are identified then it is closest to the gene. |
| TRANSFAC<sup>23</sup> | Curated Transcription Factor Targets dataset based on Positional Weight Matrices (PWMs). | PWMs are used to uncharacterised bindings sequences to closet gene using matrix based search of TF binding sites. |
| CHEA<sup>24</sup> | ChIP-X Enrichment Analysis from published ChIP-X experiments. | Target calling based on 400 bp sliding window to detect overlap between peak and closest gene. |

Trans-genes were categorised according to evidence of GR regulation from datasets shown in Table 2. Genes were scored against these criteria: 1) appearing in a transcription factor database (ENCODE, TRANSFAC, CHEA); 2) identified as GR target from ChIP-seq experiment in adipocytes from Yu *et al*^20^; 3) differentially expressed in response to dexamethasone treatment in adipocytes from Yu *et al*^20^; and 4) murine homolog of human gene differentially expressed in murine RNA-seq experiments (FC > 1; p-value < 0.05)^21^. Genes were then ranked according to how well they met the criteria for GR regulation (+1 for each item matched from criteria 1-4).

### 2.5 Causal gene network reconstruction

Cis-eQTL discovery was carried out to identify genetic instruments to be used for causal inference analysis with Findr^19^ (Figure S3). An automated pipeline was established to use the secondary linkage test (P2) to calculate SNP-gene associations when supplied with a list of genes. SNP-gene associations were obtained between all SNPs within 1 Mb of the trans-gene and all other transcripts using the same tissue dataset as the trans-gene.

Associations between all SNPs and the trans-gene were extracted from the output. A primary cis-eQTL was selected for each gene, defined as the SNP-gene association with the highest Findr P2 score for the trans-gene. An alternate, independent, cis-eQTL was selected as the second strongest cis-association not in Linkage Disequilibrium (LD) with the primary cis-eQTL. LD between SNPs was calculated as the Pearson correlation coefficient between the primary cis-eQTL genotype and all other SNP genotypes. The alternate cis-eQTL was defined as the top cis-association, which was not in LD with the primary cis-eQTL (R^2^ < 0.5).

To test for pleiotropy between the selected instrument and other cis-genes, cis associations between all cis genes (± 1 Mb of the trans-gene) and the primary instrument were obtained, as detailed in supplementary information (S2: Application of independent genetic instruments for gene network reconstruction)

All genes with a valid cis-eQTL (P2 ≥ 0.75) were taken forward for causal analysis with Findr. Causal relationships were inferred between these cis-eQTL genes (A-genes) and all other transcripts expressed in the same tissue (B-genes). The input was as follows: (dg) array of eQTL genotypes A-gene in 012 format, (d) array of normalised A-gene expression levels, (dt) array of expression levels for all B-genes in the relevant tissue sorted with d appearing on top.

The output of all tests in Findr was calculated using the *pijs_gassist* function from the Findr Python package. The posterior probability of a causal interaction (P(A→B)) was calculated from the product of the alternative hypotheses from the secondary linkage test (P2) and the controlled test (P5). The controlled test (P5) is a likelihood ratio test, which can be used as a composite test with secondary linkage (P2*P5) to infer a causal A→B relationship while using a cis-eQTL, E, as an instrumental variable. P5 examines whether A and B are not associated independently with E (i.e. whether they are still coexpressed after adjusting for E), while P2 tests for a direct association between E→B. Previous work has demonstrated that most cis-eQTLs are only associated with a single gene^25^, therefore selecting cis-eQTLs specifically as an instrument allows E→B to be used as a proxy for estimating causal effects between A→B. When combined with P2, P5 can then be used to account for the comparatively few instances where E is a cis-eQTL for more than one gene, although in such cases a false positive may still occur when A and B are confounded by a common regulator^19,26^. Therefore, we also examined manually all cis-associations for selected E of interest, to account for any sources of pleiotropy that may have been missed by P5 (S2: Application of independent genetic instruments for gene network reconstruction).

This approach was undertaken for each A-gene in a given tissue in a iterative fashion. Following completion of analysis for all A-genes in a tissue, the output was converted from the default matrix format to a Pandas DataFrame. Each tissue-specific gene set of A→B pairwise interactions was filtered according to a local precision FDR threshold (Findr score) for each interaction, to correspond to a global FDR for all interactions in the tissue set.

Networks were assembled, using the network visualisation tool, Cytoscape (version 3.8.0), from FDR thresholded pairwise gene interactions previously described. These were assembled as directed networks where the A-gene acts as the parent node and the B-gene as the child node, with the posterior probability of an A→B interaction forming the network edge.

### 2.6 Setting a global FDR threshold

The Findr score for a given A→B pairwise interaction, or E→B in the case of P2 testing (Table S2), is calculated as 1 minus the probability of that interaction being a false positive. To obtain the probability of a false positive across all interactions in a gene set, this was calculated as 1 minus the mean of all local precision FDR scores for a given tissue. A Findr score cut off was then set to obtain interaction sets at 10%, 15% and 20% global FDR thresholds^27^.

### 2.7 Functional annotation and clustering for GO enrichment

Gene sets were functionally annotated using the Database for Annotation, Visualization and Integrated Discovery (DAVID)^28^. This web-based application allows for the generation of gene clusters that have been grouped in relation to an enrichment of functional terms, including but not limited to Gene Ontology (GO) terms. The strength of the gene-term interactions are measured by EASE scores, a modified Fisher’s exact test. An enrichment score for a given cluster is generated as the geometric mean of all the EASE scores within a cluster that has undergone -log transformation. For all analyses Ensembl Gene IDs were used as the input format for DAVID as opposed to universal gene symbols.

For the analyses conducted, all of the default annotation options were selected in addition to: GAD DISEASE, GO TERM BP FAT, GO TERM CC FAT, GO TERM MF FAT, PUBMED ID, REACTOME PATHWAY, BIOGRID INTERACTIONS and UP TISSUE. Gene sets were then run using DAVID and functionally enriched clusters generated using high classification stringency. Tissue-specific gene sets from STARNET RNA-seq datasets were used as background for enrichment (Table 1).

### 2.8 Transcription factor target enrichment

Lists of known transcription factor targets for both *NR3C1* and *IRF2* were obtained from ENCODE and TRANSFAC datasets respectively. These datasets were used to test for an enrichment of known transcription factor targets within novel gene sets derived from gene network targets. This was performed using Fisher’s exact test from the Python module Scipy Stats and involved the creation of a 2×2 contingency table based on a tissue-specific background consisting of all genes available in the corresponding tissue.

### 2.9 Gene network replication

Correlations between gene network targets were calculated using gene expression data from STARNET, STAGE and METSIM. Gene expression matrices were filtered to only include the target genes under investigation. The *.corr* function in Pandas was used to construct correlation matrices of corresponding Pearson correlation coefficients as absolute values.

A background gene set was constructed from the overlapping genes between the STARNET gene expression set that was used for network discovery and the corresponding gene expression set that was being used for replication. The previously described correlation analysis was then repeated using a random set of genes (the same size as the target set) selected from the background gene-set using the *.sample* function in Pandas. The Kruskal Wallis test was used in Scipy Stats to test if the targeted and randomly sampled correlations follow the same distribution. Both the targeted and random correlations were then plotted as a box plot using the Python plotting package Seaborn.

### 2.10 Gene expression clustering

Hierarchical clustering was performed on correlation values between network targets using the discovery (STARNET) gene expression data and hierarchical clustering from Scipy in Python. The leaves list that resulted from the clustering of the discovery dataset was then extracted and applied to the correlations between target genes from the corresponding replication dataset. Both sets of clustered correlation values were then plotted as opposing correlation heatmaps in Python.

## 3 Results

### 3.1 Cortisol-associated trans-genes

SNPs associated with plasma cortisol at the *SERPINA6/SERPINA1* locus have previously been linked as expression Single Nucleotide Polymorphisms (eSNPs) for *SERPINA6* in liver^7^. Using genotype and tissue-specific RNA-seq data from the STARNET cohort (Table 1) we explored the hepatic and extra-hepatic consequences of genetic variation for plasma cortisol using 73 cortisol-associated SNPs at genome wide significance (p ≤ 5×10^-8^) identified from the COR-NET GWAMA^7^. We identified 704 eQTL associations in cis and trans between plasma cortisol-associated SNPs and genes measured across all STARNET tissues, composed of 262 unique genes and 72 SNPs at a 15% FDR threshold (Table S1-S2).

The tissues with the greatest number of trans-genes were liver, subcutaneous fat and visceral abdominal fat, with a combined total of 157 trans-genes and 422 total SNP-gene associations (FDR = 15%) (Figure 1A). The vast majority of trans-eQTL associations were specific to a single tissue. A single trans-gene, the glycosyltransferase encoding gene *OGT*, was identified in both liver and visceral abdominal fat. However, as this was the only cross-tissue trans-gene identified, this suggests that the transcriptional impact of genetic variation at the *SERPINA6/ SERPINA1* locus is highly tissue-specific. The CORNET GWAMA describes 4 blocks of SNPs in LD which represent the cortisol-associated variation at the *SERPINA6/ SERPINA1* locus^7^. We observed that LD blocks 2 and 4 represent the majority of the variation across all tissues in the trans-gene sets (Figure 1B-C).

**Figure 1:**
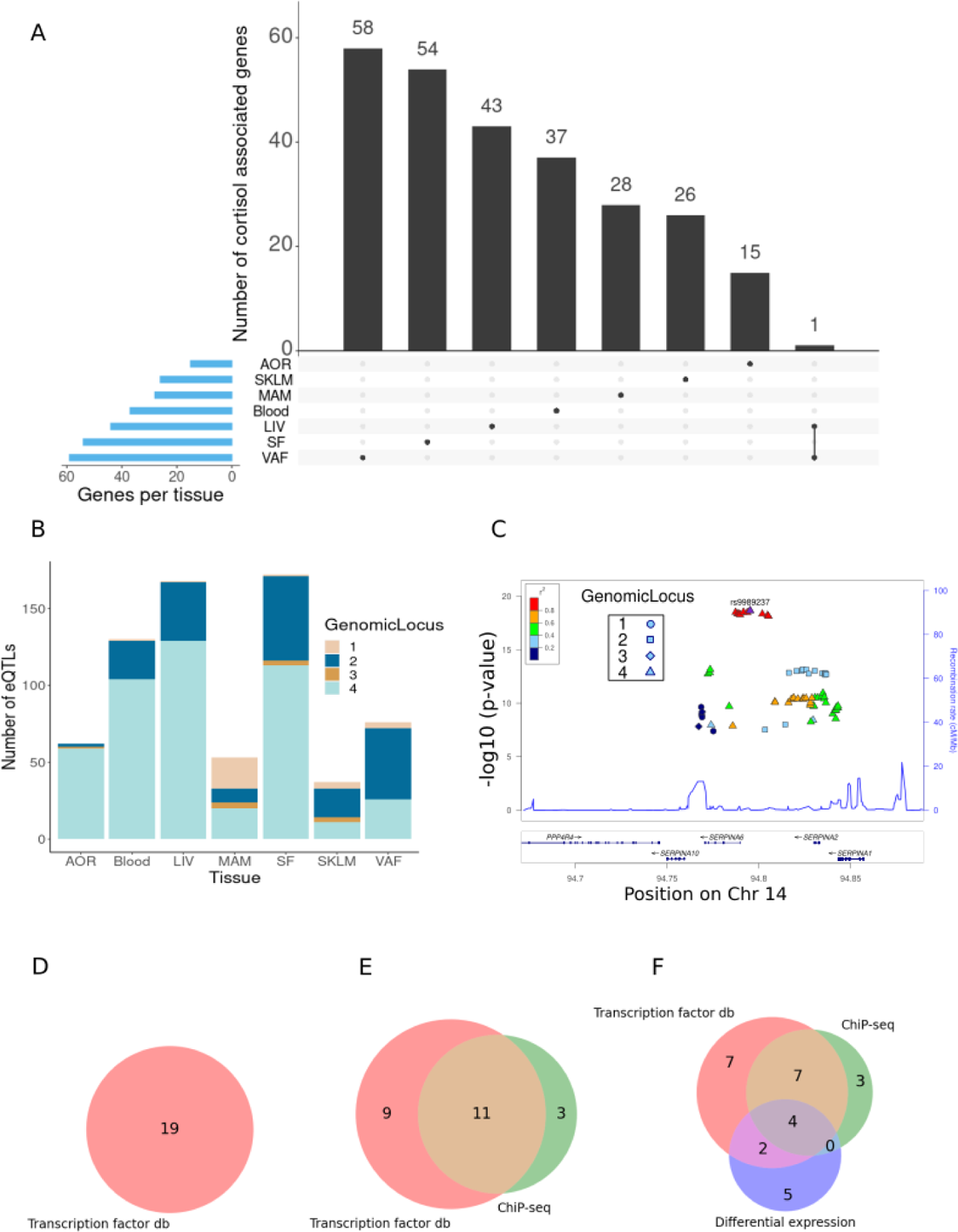
Identification of cortisol-associated trans-genes across STARNET tissues (FDR = 15%). **(A)** Upset plot showing the distribution of trans-genes across STARNET tissues, including genes shared by multiple tissues. Tissues include atherosclerotic aortic root (AOR), skeletal muscle (SKLM), internal mammary artery (MAM), blood (BLOOD), liver (LIV), subcutaneous fat (SF) and visceral abdominal fat (VAF). **(B)** Distribution of trans-eQTLs across tissues and coloured by genomic locus (LD block) of associated SNP. **(C)** LocusZoom plot^29^ showing the location of cortisol associated SNPs within defined LD blocks. **(D)** Venn diagrams where groupings represent different sources used to identify GR-linked trans-genes in liver, **(E)** visceral abdominal fat and **(F)** subcutaneous fat. These sources include transcription factor databases (db), ChiP-seq from perturbation-based experiments^20^ and differential expression of dexamethasone treated mice^21^.

### 3.2 GR regulated trans-genes associated with plasma cortisol

As GRis the primary mechanism by which cortisol influences transcription, we sought to identify a subset of cortisol-associated trans-genes that were also regulated by GR. The cortisol associated trans-genes identified in this study were compared to sets of known GR targets identified from different sources as described in Table 2. This included large projects such as ENCODE, TRANSFAC and CHEA which predict transcription factor binding targets from high throughput transcription factor binding assays. We also included predicted GR targets from perturbation-based experiments in specific tissues. ChIP-seq and microarray analysis has been used to identify 274 glucocorticoid-regulated genes in 3TS-L1 adipocytes, a murine derived cell line^20^. In addition RNA-seq data in subcutaneous fat from adrenalectomised mice treated with dexamethasone, a GR agonist, has been used to identify genes that are differentially expressed.

The greatest number of unique cortisol-associated trans-genes were identified in liver (n=43), subcutaneous fat (n=54) and visceral abdominal fat (n=59) at a 15% FDR threshold. The involvement of these tissues in glucocorticoid signaling and physiological effects has been well documented in the literature^30–33^, therefore the identification of GR-regulated trans-genes was restricted to these tissues. Comparisons of genes identified as glucocorticoid-regulated in 3T3-L1 adipocytes were only made with subcutaneous and visceral adipose trans-genes. Likewise, as the murine RNA-seq experiments were restricted to subcutaneous adipose, only subcutaneous adipose trans-genes were compared to these differentially expressed genes.

In the liver trans-gene set, 19/43 genes were identified that were present in either the ENCODE, TRANSFAC or CHEA datasets (FDR = 15%) (Figure 1C, Table S3). This includes *SERPINA6* which is cis-associated with genetic variation for plasma cortisol, as described previously^7^. One gene, *CPEB2*, was identified in more than one dataset and was present in both ENCODE and CHEA. *CPEB2* (posterior probability = 0.89) is a regulator of translation, splice variants of which have been linked to cancer metastasis^34^.

Of the cortisol-associated trans-genes identified in subcutaneous adipose (FDR = 15%), 28/54 genes were present in either a transcription factor dataset or identified from the adipose-specific perturbation datasets (Figure 1E, Table S4). There were 13 genes that had been identified as GR targets from both high-throughput transcription factor binding assays and adipose-specific experiments. These include *RNF13* which encodes IRE1α-interacting protein which plays an important role in the endoplasmic reticulum (ER) stress response through regulation of IRE1α, a critical sensor of unfolded proteins^35^. Also *IRF2*, encoding the transcription factor Interferon Regulatory Factor 2 which plays an important role as a repressor of *IRF1* which in turn is involved in the interferon-mediated immune response^36^. Furthermore, *IRF1* has previously been identified as a marker for glucocorticoid sensitivity in peripheral blood^37^.

Visceral adipose tissue had the largest number of cortisol-associated trans-genes. 21/59 of these genes had some evidence of being targets of GR (Figure 1D, Table S5). There were 5 genes that had been identified as GR targets from both high throughput transcription factor binding assays and adipose specific experiments. These include *CD163* and *LUC7L3*. *CD163* is a haemoglobin scavenger protein that is expressed in macrophages and involved in the clearance of hemoglobin/haptoglobin complexes which may play a role in protection from oxidative damage. It also plays a role in activating macrophages as part of the inflammatory response^38^. *LUC7L3*, also known as CROP, encodes a protein that is involved in alternative splicing and is associated with human heart failure^39^. It has also been shown to play a role in the inhibition of hepatitis B replication^40^.

### 3.3 Reconstruction of cortisol-associated gene networks

Having identified cortisol-associated trans-genes that are regulated by GR, causal estimates were obtained for pairwise relationships between GR-regulated trans-genes and all other genes within the given tissue. This was carried out for all GR-regulated trans-genes in liver, subcutaneous fat and visceral abdominal fat with a valid cis-eQTL instrument (12, 19, and 7 genes respectively) (Table S6). A 10% global FDR threshold was then imposed for each gene set (Table 3). Primary networks were obtained by filtering to include only GR trans-genes with a minimum of 4 target genes at the global FDR threshold.

**Table 3:** Number of network targets following FDR filtering. Total targets includes all pairwise interactions at given threshold and network regulators correspond to trans-genes with at least 4 network targets at the given FDR threshold. Inclusive of network regulators present at both 10% and 15% thresholds.

| Tissue | FDR threshold | Total targets | Network regulator | Regulator targets |
| --- | --- | --- | --- | --- |
| liver | 15% | 197 | *CPEB2* | 190 |
|  | 10% | 48 | *CPEB2* | 44 |
| subcutaneous fat | 15% | 1701 | *RNF13* | 416 |
|  |  |  | *IRF2* | 247 |
|  |  |  | *PBX2* | 883 |
|  | 10% | 486 | *RNF13* | 215 |
|  |  |  | *IRF2* | 128 |
|  |  |  | *PBX2* | 138 |
| visceral abdominal fat | 15% | 396 | *CD163* | 378 |
|  |  |  | *LUC7L3* | 15 |
|  | 10% | 17 | *CD163* | 4 |
|  |  |  | *LUC7L3* | 11 |

In liver, we identified a single gene network driven by *CPEB2*, which was found to be trans-associated with the cortisol associated SNP rs4905194 (Figure 2A). This network contained 48 causal interactions driven by *CPEB2* at a 10% FDR threshold (Figure 2D, Table S8). It is notable that *CPEB2* appears as the only network regulator in liver considering it was also the cortisol-associated trans-gene with the strongest links to GR regulation from the liver trans-gene set. A detailed description of the *CPEB2* network and all other networks identified can be found in the supplementary information (S1: Detailed networks description).

**Figure 2:**
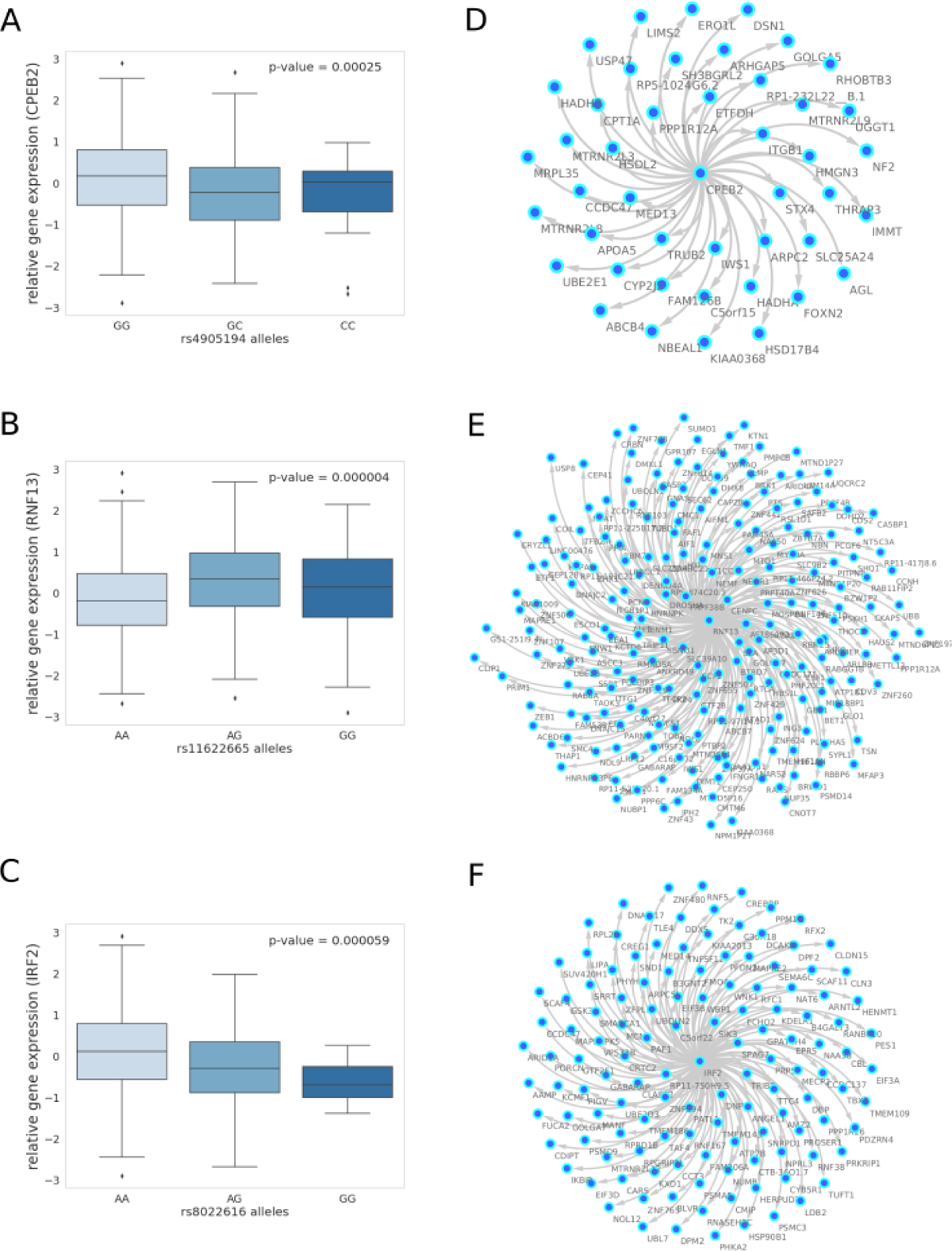
10% FDR gene networks in STARNET across different tissues. **(A)** Gene expression boxplot in liver showing trans-association with cortisol-linked SNP rs4905194 and *CPEB2*, **(B)** in subcutaneous fat between rs11622665 and *RNF13* and **(C)** rs8022616 and *IRF2* (p-value obtained from Kruskal Wallis test statistic). Box shows quarterlies of the dataset, with whiskers indicating the upper and lower variability of the distribution. **(D)** Causal gene network reconstructed from pairwise interactions from GR-regulated trans-genes against all other genes in the corresponding tissue for *CPEB2*, **(E)** *RNF13* and **(F)** *IRF2*. Edges represent Bayesian posterior probabilities of pairwise interaction between genes (nodes) exceeding 10% global FDR. Arrow indicates direction of regulation and interactions were only retained where parent node had at least 4 targets.

In subcutaneous fat, 2 major sub-networks were identified under the regulation of the genes *RNF13* and *IRF2*. This includes a total of 343 causal relationships across both sub-networks, including two genes shared by both sub-networks. *RNF13* was found to be trans-associated with the cortisol-associated SNP rs11622665 (Figure 2B) and represents the largest subcutaneous fat sub-network with 215 gene targets at a 10% FDR threshold (Figure 2E, Table S9).

The transcription factor *IRF2*, which was associated with the cortisol-linked SNP rs8022616 (Figure 2C), was found to putatively regulate a network of 128 genes (FDR = 10%) (Figure 2F). Some notable targets of *IRF2* include *LDB2* (Posterior probability = 0.94) and *LIPA* (Posterior probability = 0.91). GWAS suggests functions for *LIPA* related to CAD and ischaemic cardiomyopathy^41^, whilst *LDB2* has been demonstrated to be involved in the development of atherosclerosis^42^. Additionally, cortisol has been shown to induce a 5-fold reduction in *LDB2* expression in adipocytes^43^.

Predicted *IRF2* transcription factor targets have been previously described as part of the TRANSFAC dataset. We examined the overlap between predicted *IRF2* targets in TRANSFAC and gene targets within the *IRF2* causal networks we identified in subcutaneous fat. A true network of *IRF2* targets would be expected to show an enrichment of predicted *IRF2*. Using Fisher’s exact test in data from subcutaneous fat, at a 10% FDR threshold, the *IRF2* network had 128 target genes, 35 of which were also predicted *IRF2* targets (p = 0.08); at a 15% FDR threshold, 104/247 causal targets were also predicted targets of *IRF2* in TRANSFAC (p = 0.005). Decreasing the global FDR beyond this threshold increased the number of TRANSFAC targets within the pool of causal targets, however at a lower enrichment (p = 0.046) (Table 4).

**Table 4:**
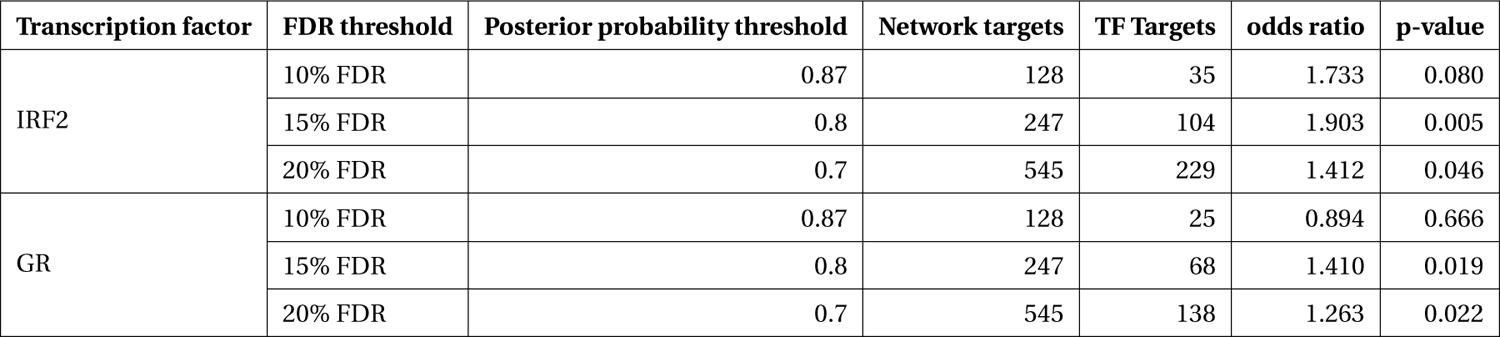
Transcription factor enrichment within gene network targets. Fisher’s exact test for *IRF2* targets from TRANSFAC predicted targets and GR targets from ENCODE. STARNET subcutaneous fat genes used as background for enrichment.

In addition to examining the prevalence of *IRF2* targets within the *IRF2* causal network, we investigated the overlap between network genes that are also regulated by GR. We observed an enrichment of ENCODE GR targets at 15 and 20% FDR thresholds (p ≤ 0.05) including 68 and 138 GR targets respectively. No GR enrichment was observed in either CHEA or TRANSFAC datasets for *IRF2* networks.

### 3.4 Co-expression of cortisol network targets in independent datasets

Causal gene networks represent coordinated changes in gene expression in response to changes in expression of network regulators. Therefore, it is possible to examine if these changes in gene expression are present in independent datasets using gene expression data alone. We used RNA-seq and microarray data from METSIM and STAGE datasets, respectively, to compare patterns in gene expression within causal networks predicted from STARNET. As METSIM only contains gene expression data for subcutaneous fat, analysis was restricted to the causal networks identified in STARNET subcutaneous fat.

Absolute correlation coefficients between the targets of the previously described network regulators were calculated and their distributions were compared to distributions of random sets of genes selected from the replication gene expression data, the same size as the corresponding target gene set. The difference between targeted and random distributions was formalised using the Kruskal Wallis test for each sub-network (Table 5).

**Table 5:** Correlations between network targets within replication datasets. Kruskal Wallis test calculated for distribution of correlations between network targets compared to correlations within random gene set of same size.

| Replication dataset | Tissue | Network regulator | p-value | No. target genes |
| --- | --- | --- | --- | --- |
| METSIM | Subcutaneous fat | *IRF2* | $< 1.0 \times 10^{-300}$ | 128 |
| | | *RNF13* | $2.3 \times 10^{-7}$ | 215 |
| STAGE | Liver | *CPEB2* | $8.2 \times 10^{-32}$ | 44 |
| | Subcutaneous fat | *IRF2* | $8.3 \times 10^{-86}$ | 128 |
| | | *RNF13* | $< 1.0 \times 10^{-300}$ | 215 |
| | Visceral abdominal fat | *CD163* | $2.6 \times 10^{-3}$ | 4 |
| | | *LUC7L3* | $4.4 \times 10^{-1}$ | 11 |

In liver, correlations between network targets of the single sub-network under the regulation of *CPEB2* were observed in STARNET and STAGE. Hierarchical clustering within STARNET liver also revealed clustering of correlated genes which were retained when the clustered gene order was then applied to STAGE liver (Figure 3A). Correlations between the 44 *CPEB2* target genes in STAGE-liver were stronger than their random counterparts (p = 8.2×10^-32^), with this shift also being observed in STARNET liver (p = 2.32×10^-197^) (Figure 3D).

**Figure 3:**
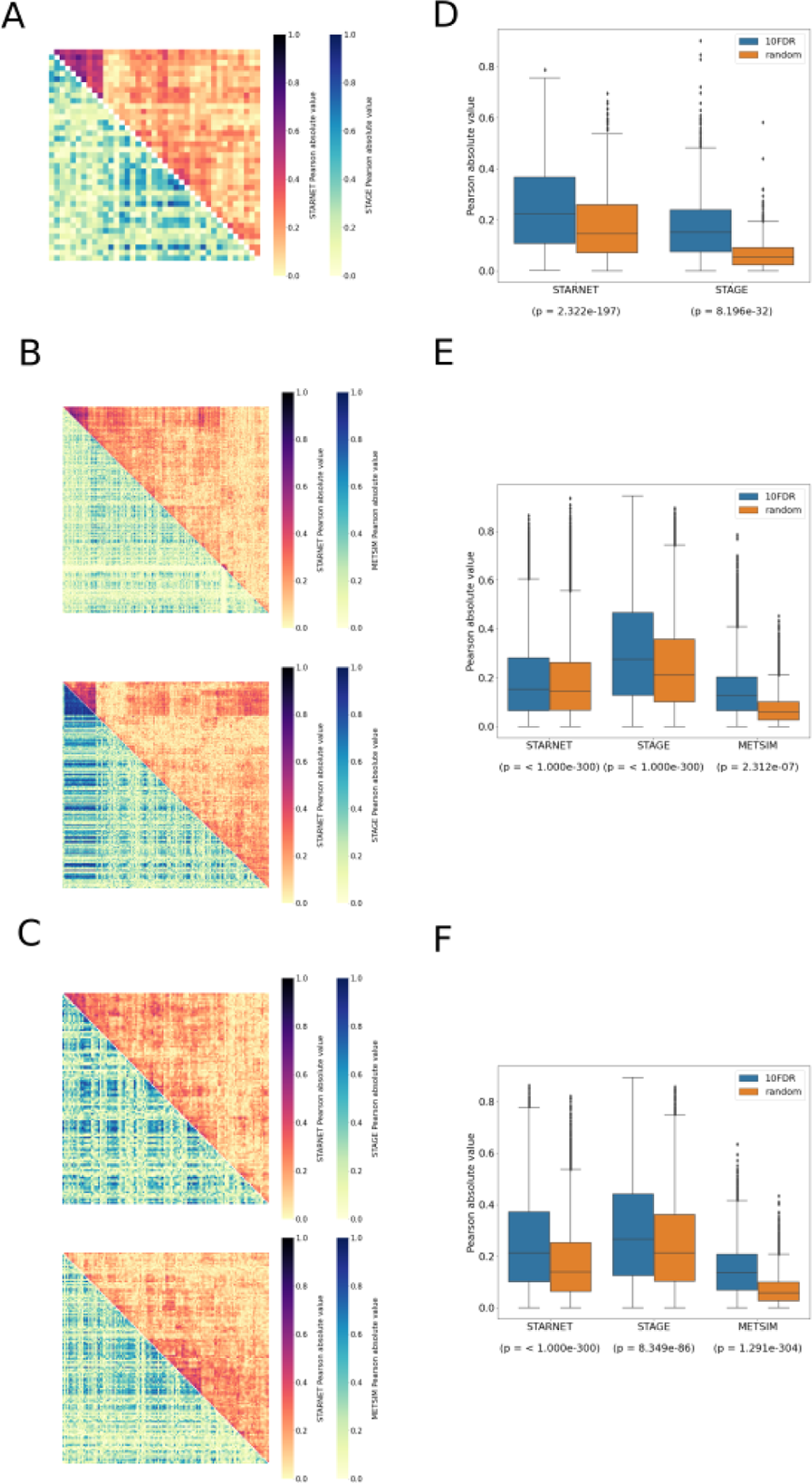
Replication of cortisol-associated gene networks in independent datasets. **(A)** Correlation heatmap showing pairwise Pearson correlations between *CPEB2*, **(B)** *IRF2* and **(C)** *RNF13* network targets. Hierarchical clustering of genes in STARNET (discovery) was applied to the same genes within replication datasets. **(D)** Correlations between network targets in discovery vs replication datasets for *CPEB2*, **(E)** *IRF2* and **(F)** *RNF13* networks. Kruskal Wallis test calculated for distribution of correlations between network targets compared to correlations within random gene set of same size.

In subcutaneous fat, correlations were observed between the network targets of *RNF13* and *IRF2*, and hierarchical clustering patterns from STARNET were applied to the replication datasets of STAGE and METSIM (Figure 3B-C). For *RNF13*, similar patterns of co-expression were observed in STAGE subcutaneous fat following clustering, however this was not the case in the METSIM dataset (Figure 3B). Despite this, *RNF13* targets appeared more highly correlated than their randomly selected counterparts in STARNET (p < 1.0×10^-300^), STAGE (p < 1.0×10^-300^) and to a lesser extent in METSIM (p = 2.3×10^-7^) (Figure 3E).

In subcutaneous fat, patterns of co-expression between *IRF2* targets were conserved most prominently in METSIM, however co-expression was less strongly correlated compared with *RNF13* targets (Figure 3C). *IRF2* subcutaneous fat sub-network targets were more strongly correlated than their random counterparts in STARNET (p < 1.0×10^-300^), STAGE (p = 8.35×10^-86^) and METSIM (p < 1.0×10^-300^) (Figure 3F)

## 4 Discussion

In this study we have characterised the impact that genetic variation for plasma cortisol has upon tissue-specific gene expression. We showed that cortisol-linked genetic variants at the *SERPINA6/ SERPINA1* locus mediate changes in gene expression in trans across multiple tissues, in addition to the cis-associations in liver which have been described previously^7^. We have scrutinised these trans-associations to identify a subset of genes that are regulated by glucocorticoids, and in turn regulate downstream transcriptional networks, thus providing a deeper understanding of the transcriptional landscape driven by cortisol-linked genetic variation that may underpin the progression to cardiovascular disease.

CBG, as encoded by *SERPINA6*, is responsible for binding cortisol in the blood. It has remained uncertain whether variation in CBG impacts the availability of cortisol within tissues, since any resulting change in free cortisol concentrations would be expected to be adjusted by negative feedback of the HPA axis^44^. However, deleterious mutations in CBG are associated with dysfunction in animals and humans, suggesting an impact of CBG on cortisol signaling^44^. Our major finding that downstream transcriptomic changes in extra-hepatic tissues are associated with genetic variation at the *SERPINA6* locus lends strong support to the hypothesis that CBG influences tissue delivery of cortisol and modulates glucocorticoid-induced changes in gene expression.

For the STARNET study, whole blood samples were taken pre-operatively and all other tissues including liver were taken during CABG surgery. In addition to any rise in cortisol due to anxiety and disturbed sleep in anticipation of surgery, the human stress response to surgery has been well characterised and results in stimulation of the HPA axis leading to high levels of cortisol in the blood both during and post-surgery^45^. Surgery is also associated with a very rapid fall in CBG production. Therefore, it is uncertain if cortisol-associated gene expression patterns observed in STARNET would also be observed in an un-stressed healthy population. It may be that CBG influences the dynamic range of alterations in free plasma cortisol during stress rather than affecting delivery of cortisol to tissues in unstressed conditions. However, considering that coexpression of the network targets was reproducible within independent samples from the METSIM study, obtained under non-surgical conditions, this suggests that the cortisol-associated networks we inferred from STARNET do operate also in unstressed conditions.

The tissues with the greatest number of trans-genes identified were liver and both subcutaneous and visceral abdominal fat, all tissues known to play a role in glucocorticoid biology. In liver, glucocorticoids have extensive effects on glucose and fatty acid metabolism^30,31^, while in adipose tissue glucocorticoids regulate lipogenesis and lipid turnover^32,33^. Skeletal muscle is also a major target of glucocorticoids, where they modulate protein and glucose metabolism^46^. A lack of available data for identifying tissue-specific GR targets in other tissues means that potential GR targets may have been missed in tissues outside of liver and adipose.

We identified a subset of GR-responsive genes in liver, subcutaneous fat and visceral adipose fat. However, we did not observe a statistical enrichment of GR-regulated genes in any of these trans-gene sets. This does not negate the identification of GR targets that are associated with plasma cortisol, but it may imply that there are some effects of cortisol-linked genetic variation that are mediated by mechanisms other than directly by GR, either through secondary regulation by GR-regulated genes or through the alternative mineralocorticoid receptor. Indeed, some of the genes with higher levels of evidence for GR regulation also demonstrated regulation of transcription networks e.g. *CPEB2*, *IRF2*, *RNF13*. This supports our strategy of setting a relatively lenient FDR threshold (15%) and then filtering to identify cortisol-associated trans-genes with prior evidence of GR regulation.

We identified causal gene networks in liver, subcutaneous fat and visceral abdominal fat where cortisol-associated trans-genes act as regulators of sub-networks within overarching tissue specific networks. Pairwise causal relationships were established between network regulators and downstream targets using cis-eQTLs as genetic instruments. This approach has the benefit of generating directed relationships between a regulator and target while accounting for any unobserved confounding. However, a drawback of this approach is that we are limited by only being able to examine GR regulated trans-genes with valid cis-eQTLs. This means that there could be valid cortisol responsive networks regulated by GR trans-genes which we were unable to predict due to lack of a corresponding instrument.

*IRF2* stands out as a network regulator of particular interest. There is strong evidence of GR regulation, where *IRF2* has been identified as a GR target from published dexamethasone treated adipocyte ChIP-seq experiments^20^ and as a putative GR target within ENCODE. It is robustly associated with its corresponding cis-eQTL instrument and there is an enrichment of *IRF2* targets within our predicted *IRF2* regulated causal network. Additionally, we show evidence of regulation by glucocorticoids within the targets of *IRF2*, potentially suggesting evidence of a feed-forward loop motif^47^. Interestingly the genotype for rs8022616, the cortisol associated SNP linked to *IRF2* expression in subcutaneous fat, is associated with a decrease in *IRF2* expression. Previous evidence suggests that interferon signaling is inhibited by glucocorticoids^48,49^.

In conclusion, we have linked genetic variation for plasma cortisol to changes in gene expression across the genome, beyond that which has been previously described at the *SERPINA6/ SERPINA1* locus^7^ and extending to adipose tissue as well as liver. Furthermore, we have shown that a subset of these trans-genes are driven by GR and in turn drive transcriptional networks across different tissues. These networks have been found to be robust and their network targets appear co-expressed within independent gene expression datasets of the same tissue. Further study of these networks and their downstream targets could be used to enhance our mechanistic understanding of the pathways linking cortisol with complex disease as observed in observational studies.

## 5 Author contributions

SB, TM and BW contributed to the conception and design of this research. SB conducted all formal analyses and visualisations and wrote the manuscript, supervised by TM, BW and RA. LW and TM developed and supported use of and interpretation of outputs from the software Findr. AC contributed to data analysis and interpretation for the CORNET consortium. RM conducted the experiments and contributed to data analysis of dexamethasone-treated mice. AR and JLMB provided access to and contributed to interpretation of data from the STARNET cohort. All authors reviewed the manuscript and approved the submitted version.

## 7 Data and code availability

All code used in the analyses presented in this study are available at the following repository: https://github.com/sbankier/cortisol_networks/tree/main.Data from the Stockholm Tartu Atherosclerosis Reverse Networks Engineering Task study (STARNET) are available through a database of Genotypes and Phenotypes (dbGaP) application (accession no. phs001203.v2.p1). Gene expression data from The Stockholm Atherosclerosis Gene Expression study (STAGE) and the Metabolic Syndrome in Man study (METSIM) are available publicly at GEO (accession no. GSE70353 and GSE40231, respectively). The summary statistics from the CORNET GWAMA are available at Edinburgh DataShare (https://datashare.ed.ac.uk/handle/10283/3836).

## Supplementary material

### S1 Detailed networks description

#### S1.1 Liver

A trans-association was identified between cortisol-associated SNP rs4905194 and *CPEB2* in STARNET-liver (Figure 2A). Following cis-eQTL discovery for *CPEB2*, a SNP peak was identified upstream of the *CPEB2* TSS represented by the instrument rs62410848, which was used as an instrumental variable for network reconstruction (Figure S6).

48 causal interactions were obtained at a global 10% FDR threshold (Posterior probability ≥ 0.855) (Table SS8). When filtering to a minimum of 4 targets, the only GR-regulated trans-gene that remained was *CPEB2* (Figure 2E). Notably, this was also the trans-gene that appeared in the most GR target datasets, forming a network with 44 target genes. Functional enrichment was performed using DAVID for all *CPEB2* target genes (Table SS7). The strongest cluster was related to fatty acid beta oxidation and lipid metabolism, including 5 genes related to GO:0006635 - fatty acid beta-oxidation (adj p-value = 0.002). Other enrichments stem from 8 genes related to acquired immunodeficiency syndrome and disease progression (adj p-value = 0.003).

The strongest causal relationship within this network was between *CPEB2* and the gene *HADHA* (Posterior Probability = 0.99), responsible for encoding the alpha subunit of the mitochondrial trifunctional protein^50^. Mutations affecting this protein have been linked to long-chain 3-hydroxyacyl-CoA dehydrogenase (LCHAD) deficiency, which affects the ability to metabolise fatty acids in the liver^51^. These mutations have also been linked to maternal acute fatty liver during pregnancy^52^.

#### S1.2 Subcutaneous fat

In STARNET subcutaneous fat, 486 causal relationships were detected at a 10% FDR threshold (Posterior probability = 0.87), which is the most out of all tissues examined (Table S9). When filtering to exclude trans-genes with less than 4 targets at this threshold, 2 major sub-networks are represented under the regulation of the genes *RNF13* and *IRF2*. This includes a total of 343 causal relationships across both sub-networks, including two genes shared by both sub-networks.

*RNF13* was found to be trans-associated with the cortisol-linked SNP rs11622665 (Figure 2E). A cis-eQTL peak of SNPs associated with *RNF13* in STARNET subcutaneous fat was identified upstream of the *RNF13* TSS, represented by the lead SNP rs9853321 which was used as a causal instrument in the reconstruction of the causal network driven by *RNF13* (Figure S6).

*RNF13* represents the largest subcutaneous fat sub-network with 215 gene targets at a 10% FDR threshold (Figure 2F). The strongest functional enrichment term for *RNF13* targets is related to Poly(A) RNA binding, where 33 targets are included for this term, GO:0044822 poly(A) RNA binding (adj p-value = 0.01), and 39 targets are included for RNA binding, GO:0003723 RNA binding (adj p-value = 0.04). Other notable terms include 23 genes related to Zinc finger motifs (adj p-value = 0.05).

*IRF2* was found to be associated with the cortisol-linked SNP rs8022616 (Figure 2C). Cis-eQTL discovery revealed associations between rs34985265 and *IRF2* expression in subcutaneous fat to obtain an instrument that could be used for causal network reconstruction (Figure S6).

The *IRF2* sub-network contains 128 targets at a 10% FDR threshold (Figure 2D). Following functional enrichment of *IRF2* targets, the strongest enrichment term included 19 genes related to Poly(A) RNA binding (p-value = 0.009), however this association was not retained following multiple testing correction. Some notable targets of *IRF2* include *LDB2* (Posterior probability = 0.94) and *LIPA* (Posterior probability = 0.91). GWAS suggests functions for *LIPA* related to CAD and ischaemic cardiomyopathy and *LDB2* has been demonstrated to be involved in the development of atherosclerosis^42^. Additionally, cortisol has been shown to induce a 5-fold reduction in *LDB2* expression in adipocytes^43^.

An additional subcutaneous fat gene network was identified for the transcription factor *PBX2* containing 138 targets at a 10% FDR threshold. However, the cis-eQTL instrument that was used to reconstruct this network was found to be associated with many other genes at the *PBX2* locus, which include causal targets within the *PBX2* network. This indicates that *PBX2* is not independently linked to this instrument and the *PBX2* causal network could be driven by a cis-gene other than *PBX2*.

#### S1.3 Visceral abdominal fat

In STARNET visceral abdominal fat, trans-associations were identified for the genes *CD163* and *LUC7L3* with the same cortisol associated SNP rs2005945 (Figure S5A-B). Although STARNET-visceral abdominal fat contained the largest number of trans-associations with cortisol SNPs, the fewest causal relationships were detected in this tissue at 10% FDR (Table S10). Two small sub-networks were detected, regulated by the genes *LUC7L3* and *CD163* composed of eleven and four targets (Figure S5C). Interestingly, when the FDR threshold is reduced to 15% the sub-network for *CD163* is expanded to include 378 targets, a much more dramatic expansion compared to reducing the threshold to 15% FDR with other regulators. The networks for *CD163* and *LUC7L3* were identified using the cis-eQTLs rs73059776 and rs6504682, respectively (Figure S5D). Due to the small size of the 10% FDR networks, functional enrichment and clustering was not carried out for either of the networks identified in visceral abdominal fat.

### S2 Application of independent genetic instruments for gene network reconstruction

To study the impact of instrument selection on the reconstruction of causal networks we examined the distribution of local cis-eQTLs for each of the GR-regulated trans-genes that was found to regulate a network. Primary instruments were selected as the strongest cis-eQTL within a 1 Mb window of the associated gene, as determined by secondary linkage test posterior probability. However, the landscape of gene expression-linked genetic variation can involve several loci associated with the expression of the same gene to differing degrees. In addition to selecting a primary cis-eQTL as an instrument, alternate independent instruments were also identified. These were defined as the second strongest cis SNP-gene association which was not in LD with the primary instrument (R^2^ < 0.5) (Figure S6).

Causal relationships in STARNET liver were defined by a GR-regulated network under the regulation of *CPEB2* (FDR = 10%). The genetic instrument used to construct this network, rs62410848 (posterior probability = 0.90), is the strongest cis-eQTL for *CPEB2*, located less than 100 Kb upstream of the *CPEB2* locus. An independent peak was identified 400 Kb upstream of *CPEB2*, represented by rs6847363 as the top cis-association in this region (posterior probability = 0.48). As this independent instrument fell below the required threshold (posterior probability ≥ 0.75), causal analysis was not carried out using rs6847363 as an instrument.

To determine the robustness of the primary instrument, we examined cis-associations with other genes within this locus (± 1 Mb). While *CPEB2* was the strongest cis-eQTL association in this region, rs62410848 was also seen to be associated with *CD38* (posterior probability = 0.85), a gene ~800 Kb downstream of *CPEB2*. Although *CD38* is not associated with any cortisol variants at the *SERPINA6/ SERPINA1* locus, it has been identified as being regulated by glucocorticoids in smooth muscle cells^53^ and has been identified as a GR target in ENCODE. However, *CD38* does not appear as a target of *CPEB2*, which suggests a low P5 score. This suggests that *CPEB2* and *CD38* are independently associated with rs62410848 and that *CPEB2* is the true network regulator in this cis region.

In STARNET subcutaneous fat, the *IRF2* sub-network was generated using the SNP rs34985265 (posterior probability = 0.94) located ~500 Kb upstream of *IRF2.* The strongest independent cis-eQTL for *IRF2*, rs2171838 (posterior probability = 0.72), is located closer to *IRF2*, ~300 Kb of the *IRF2* TSS. This association did not reach the association threshold for use as a causal instrument (posterior probability = 0.72). Examining cis-associations between rs34985265 and all genes within 1 Mb of *IRF2*, *IRF2* is the only gene to show an association with this SNP.

For *RNF13* the primary instrument, rs9853321 (posterior probability = 0.81), was located in a peak 400 Kb upstream of the *RNF13* transcription start site. The strongest independent cis-eQTL, rs62282739, is located nearly 1 Mb downstream of *RNF13* and was too weak to be taken froward for causal analysis (posterior probability = 0.70). Cis-associations for rs9853321 in this region, include an association with the gene *TM4SF1* (posterior probability = 0.93) at a higher level than the association with *RNF13*. There is some indication of a causal relationship between *RNF13* and *TM4S1* (posterior probability = 0.73), however *TM4S1* is not a target of *RNF13* at either a 10% or 15% global FDR threshold, suggesting that *TM4S1* is independently associated with rs9853321.

The third subcutaneous fat sub-network was predicted using the SNP rs35571244 as a cis-eQTL for *PBX2* (posterior probability = 0.93). This SNP is located ~800 Kb downstream of the *PBX2* transcription start site and is the strongest cis-eQTL for a peak of SNPs in this region. An alternate cis-eQTL, rs3128947 (posterior probability = 0.73), is located 500 Kb upstream of the *PBX2* transcription start site. Again, this cis-eQTL was too weak to be taken forward for causal analysis. There are 31 cis-associations between rs35571244 and genes within a 1 Mb window of *PBX2* at a 15% FDR threshold (posterior probability ≥ 0.8), of which *PBX2* is the 7th strongest association. Of these cis associations, 10 are causal targets of *PBX2* when using rs35571244 as a genetic instrument at a 15% FDR threshold and 4 are targets at a 10% FDR threshold. This suggests that these genes are not independently linked to rs35571244, which raises the possibility that these targets would be predicted from other cis-genes and not just *PBX2* specifically.

The primary instrument used to reconstruct the *CD163* sub-network in visceral abdominal fat, rs7954905 (posterior probability = 0.86), is located less than 100 Kb downstream of *CD163*. The strongest independent cis-eQTL, rs2377237 (posterior probability = 0.72), is located −500 Kb upstream of the *CD163* transcription start site, however this SNP was below the threshold for use as a causal instrument. There were 6 cis-associations at a 15% FDR threshold (posterior probability ≥ 0.78). One of these cis-genes is a target of *CD163* (posterior probability = 0.86) at a 15% FDR threshold, but no genes are targets at a 10% FDR threshold. This target gene is *CD163L1*, which is a paralog of *CD163* located downstream of of *CD163*. The peak represented by rs7954905 is located in the *CD163L1* gene body. *CD163L1* arose as a gene duplication of *CD163* and colocalises with *CD163*^54^.

The primary instrument used to reconstruct the *LUC7L3* sub-network, rs6504682 (posterior probability = 0.8), is located within the *LUC7L3* gene body. An independent cis-eQTL, rs2412130 (posterior probability = 0.7) is located in a peak ~1000 Kb upstream of *LUC7L3*. Again, this alternate cis-eQTL did not meet the threshold for use as an instrument. There is only one other cis-gene associated with rs6504682, *ANKRD40* (Findr score = 0.81), however this gene is not a target of *LUC7L3* in either the 15% or 10% FDR causal networks in visceral adipose.

### S3 Supplementary figures

**Figure S1:**
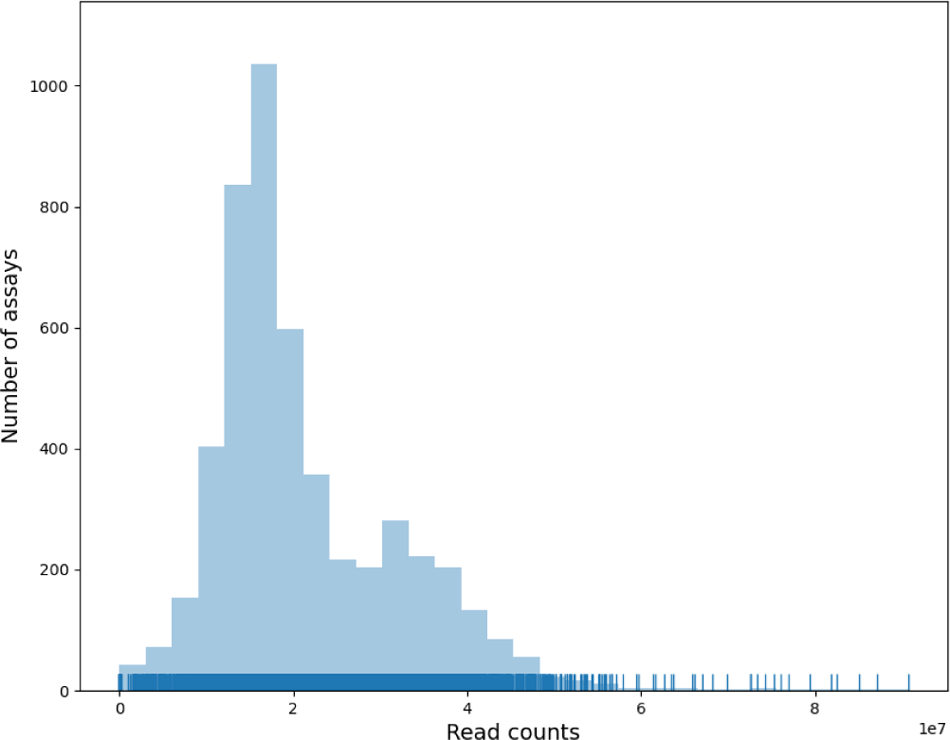
Distribution of RNA-seq read counts across all STARNET tissues.

**Figure S2:**
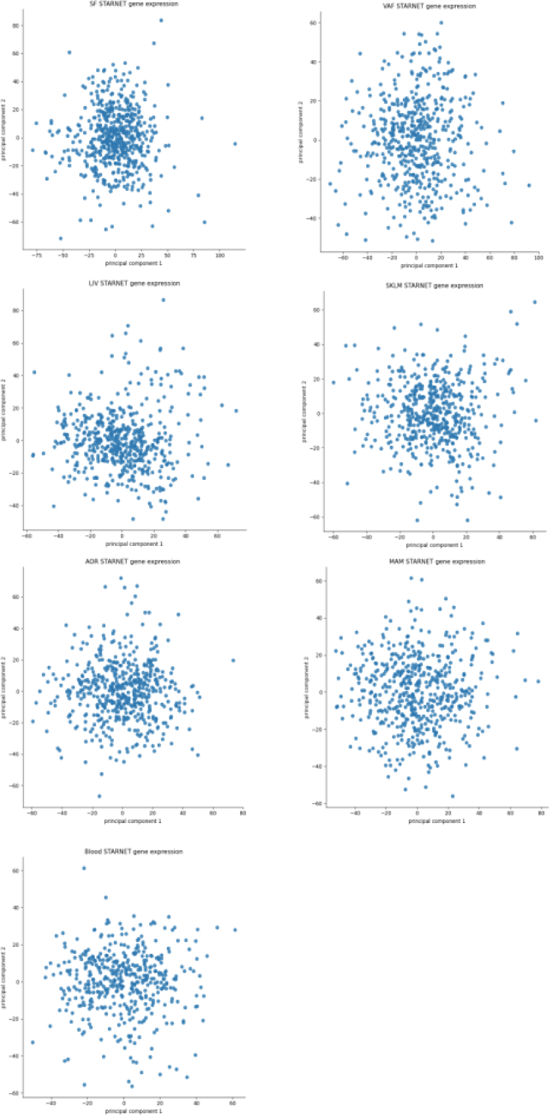
Principal component analysis of gene expression samples across all STARNET tissues.

**Figure S3:**
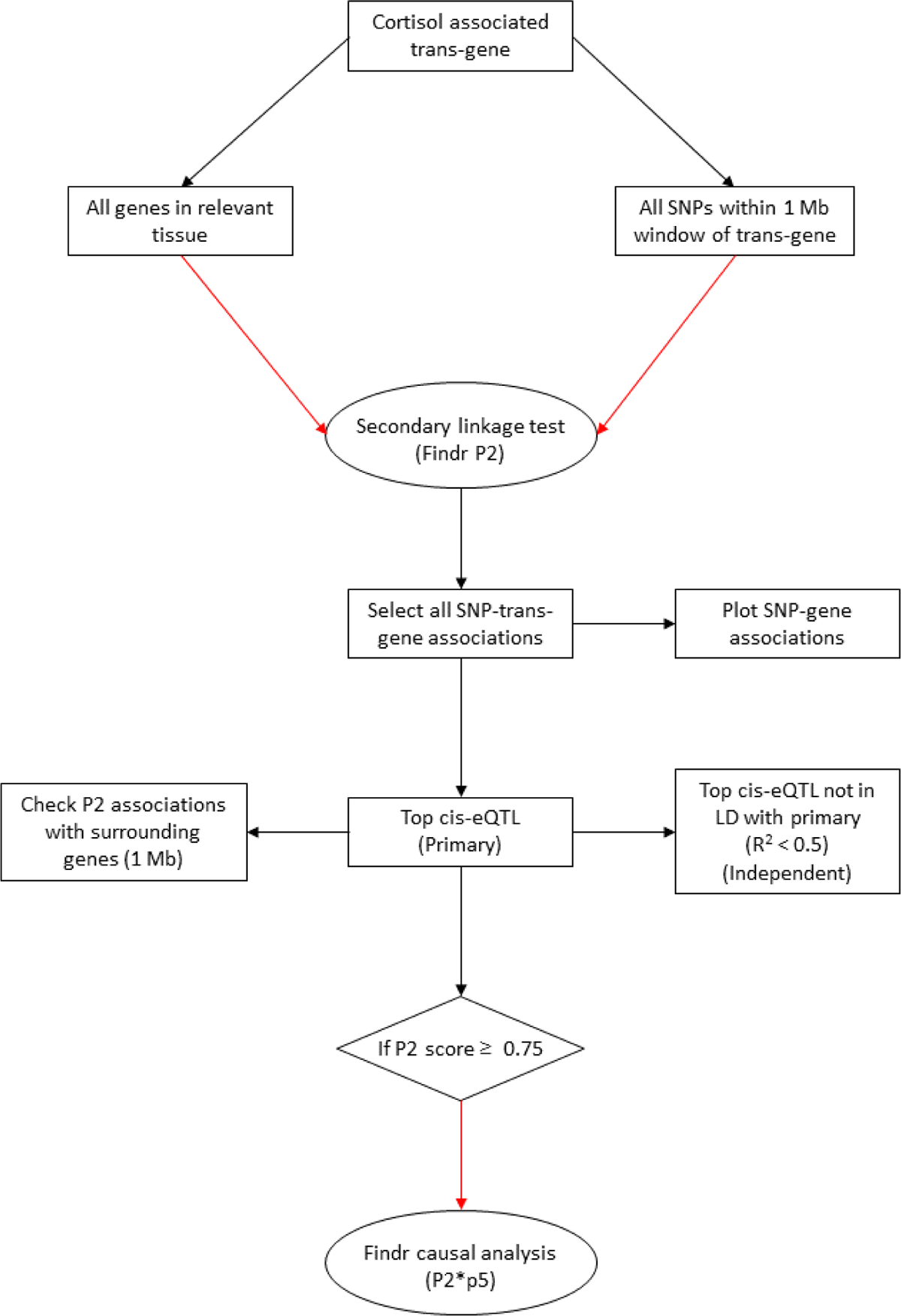
Instrument selection for causal analysis with Findr. Flowchart depicts identification of cis-eQTLs for use as genetic instruments for causal analysis with Findr.

**Figure S4:**
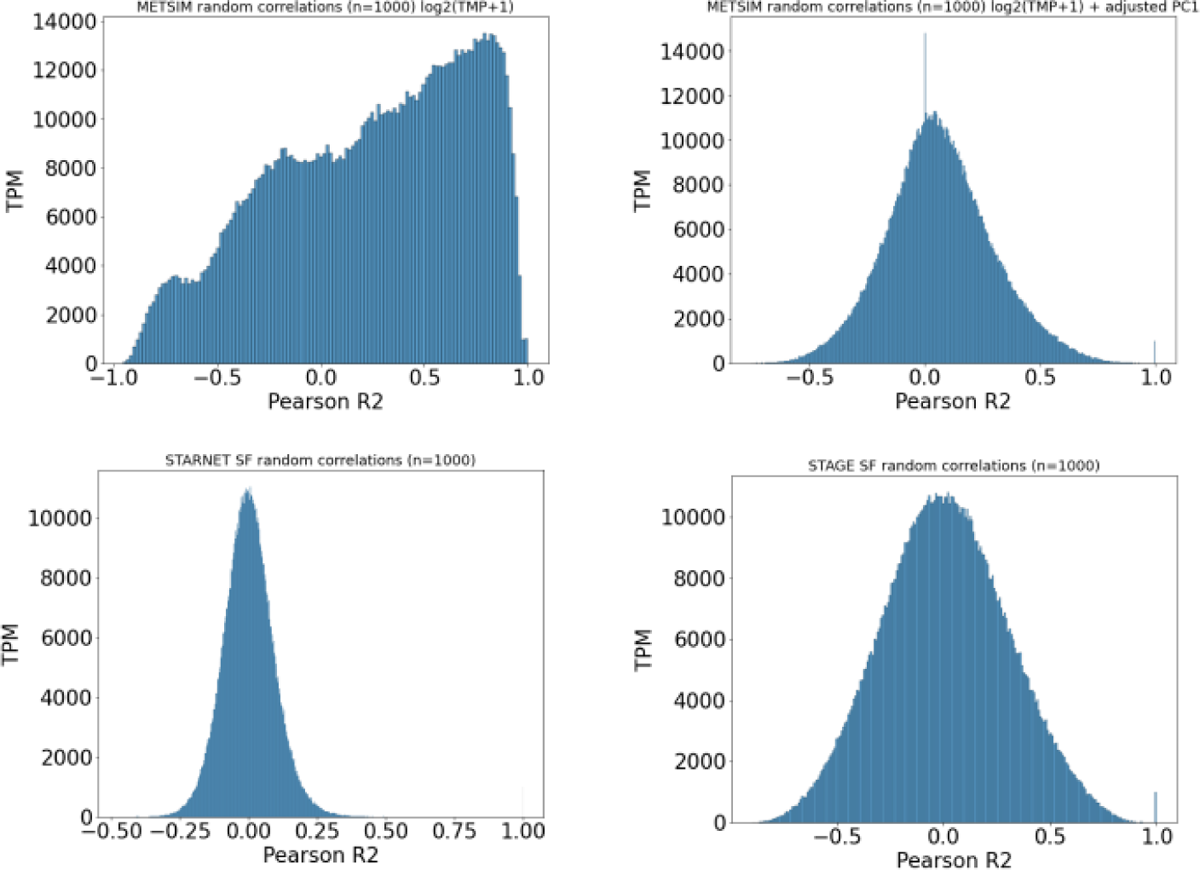
Correlations between randomly sampled genes from discovery and replication datasets both pre and post correction.

**Figure S5:**
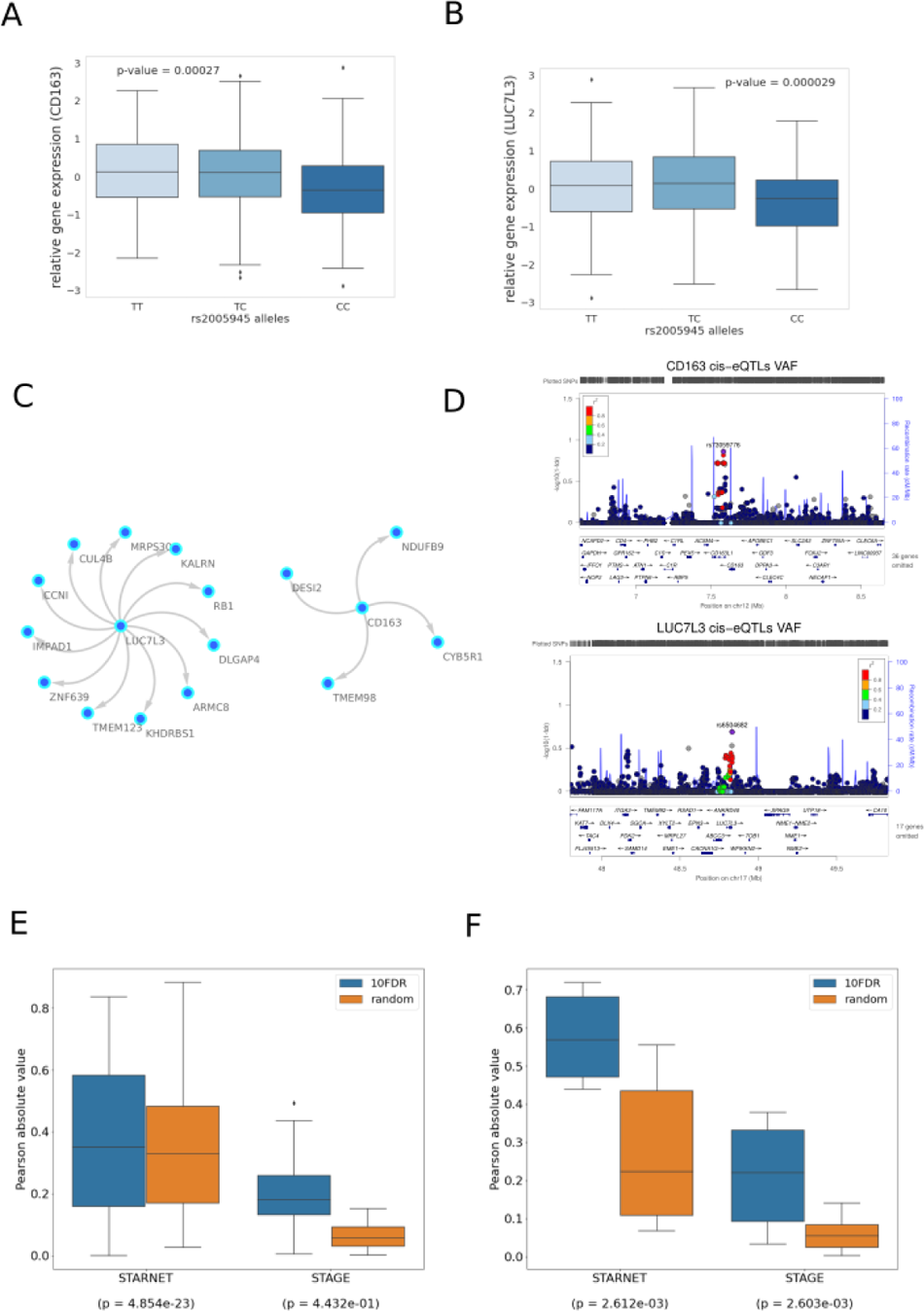
10% FDR gene network in STARNET visceral abdominal fat driven by *LUC7L3* and *CD163*. **(A)** Gene expression boxplot in STARNET visceral abdominal fat showing trans-association with cortisol-linked SNP rs20005945 and *CD163* **(B)** and *LUC7L3* (p-value obtained from Kruskal Wallis test statistic). **(C)** Causal gene network reconstructed from pairwise interactions from GR-regulated trans-genes against all other STARNET visceral abdominal fat genes. Edges represent Bayesian posterior probabilities of pairwise interaction between genes (nodes) exceeding 10% global FDR. Arrow indicates direction of regulation and interactions were only retained where parent node had at least 4 targets. **(D)** LocusZoom plot showing cis-eQTLs for *CD163* and *LUC7L3*, with lead SNP used as instrumental variable indicated in purple. Significance of association is indicated on the y axis as -log10(1-fdr), where fdr represents the local false discovery rate as estimated by Findr. **(E)** Correlations between network targets in discovery vs replication datasets for *LUC7L3* and **(F)** *CD163*. Kruskal Wallis test calculated for distribution of correlations between network targets compared to correlations within random gene set of same size.

**Figure S6:**
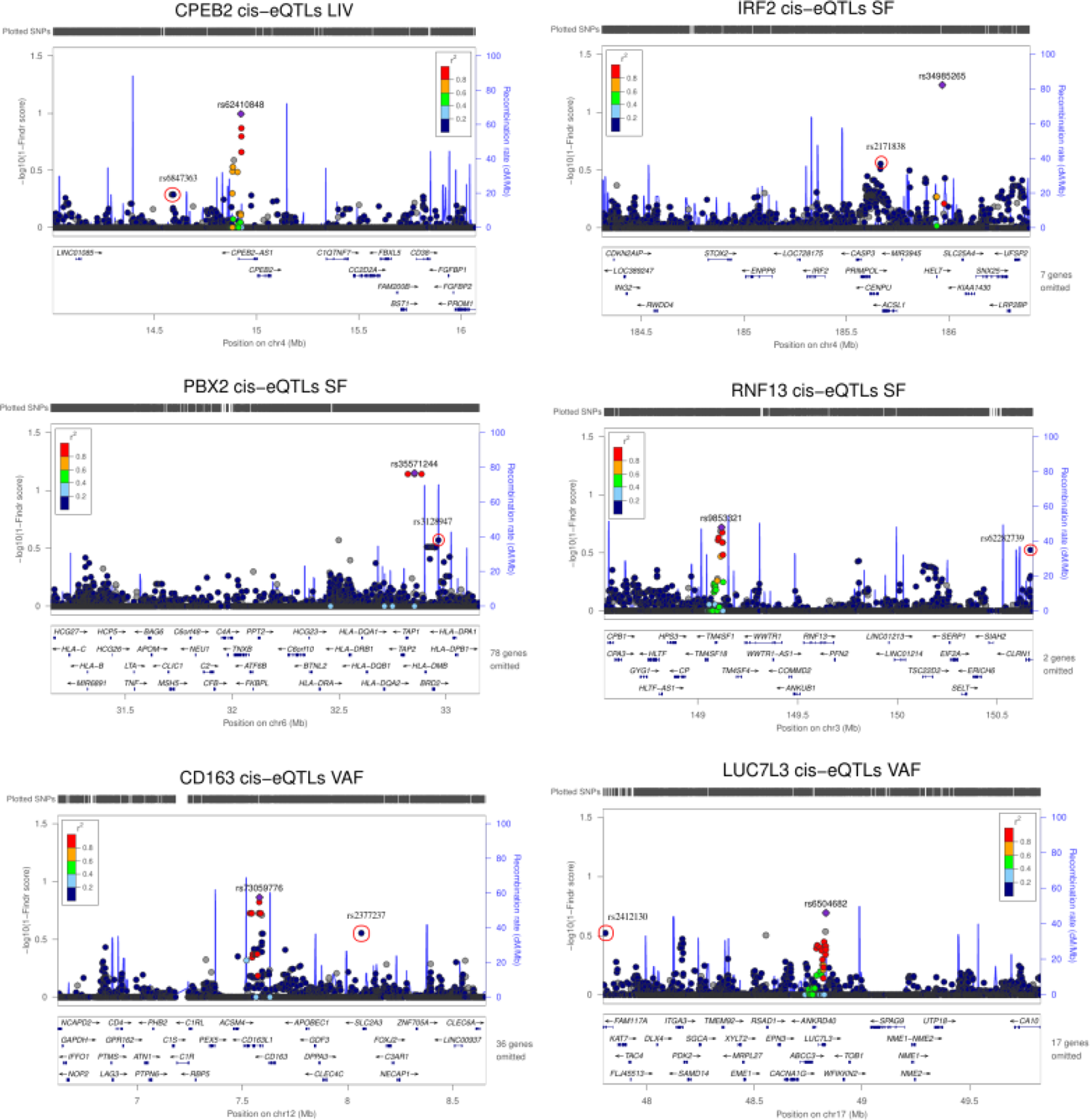
Cis-eQTL discovery for network regulators. SNP-gene associations within a 1 Mb window of the associated gene calculated using the Findr secondary linkage test (P2) and presented as 1-findr score (-log10) with LocusZoom. Lead cis-eQTL is primary instrument used for causal analysis. Red circle indicates independent (R^2^ ≤ 0.5) alternate instrument.

### S4 Supplementary tables

**Table S1:** All genes associated with variation for plasma cortisol across all STARNET tissues (FDR = 15%). Only unique associations are included with the top SNP-gene pair. Number of associations refers to the total number of cortisol associated SNPs associated with a given gene.

**Table S2:** Tissue specific local precision FDR (Findr P2 scores) used to establish FDR thresholds for trans-gene sets.

**Table S3:**Cortisol associated trans-genes from STARNET-liver (FDR = 15%) with evidence of GR regulation. Transcription factor db includes ENCODE, TRANSFAC and CHEA transcription factor datasets.

**Table S4:** Cortisol associated trans-genes from STARNET-subcutaneous fat (FDR = 15%) with evidence of GR regulation. Transcription factor db includes ENCODE, TRANSFAC and CHEA transcription factor datasets. ChIP-seq and Microarray fields are from Yu et al experiments in adipocytes^20^. Murine dex is from dexamethasone treated adrenalectomised mice^21^. Indicates genes that have been identified as GR targets from both global TF binding and perturbation experiments. Direction of effect is estimated from the Pearson correlation coefficient of the gene expression level and cortisol associated genotype.

**Table S5:** Cortisol associated trans-genes from STARNET-visceral adipose fat (FDR = 15%) with evidence ofGR regulation. Transcription factor db includes ENCODE, TRANSFAC and CHEA transcription factor datasets. ChIP-seq and Microarray fields are from Yu et al experiments in adipocytes^20^. * Indicates genes that have been identified as GR targets from both global TF binding and perturbation experiments. Direction of effect is estimated from the Pearson correlation coefficient of the gene expression level and cortisol associated genotype.

**Table S6:** GR regulated cortisol linked trans-genes (FDR = 15%) with a valid cis-eQTL for causal analysis (posterior probability ≥ 0.75).

**Table S7:** Functional enrichment of causal network targets using DAVID. Filtered to enrichment score > 1.

**Table S8:** All pairwise interactions from Findr (P2*P5) in liver at a 10% FDR threshold.

**Table S9:** All pairwise interactions from Findr (P2*P5) in subcutaneous fat at a 10% FDR threshold.

**Table S10:** All pairwise interactions from Findr (P2*P5) in visceral abdominal fat at a 10% FDR threshold.

